# Aurora A promotes chromosome congression by activating the condensin-dependent pool of KIF4A

**DOI:** 10.1101/650937

**Authors:** Elena Poser, Renaud Caous, Ulrike Gruneberg, Francis A. Barr

## Abstract

Aurora kinases create phosphorylation gradients within the spindle during prometaphase and anaphase. These locally regulate factors that promote spindle organisation, chromosome condensation and movement, and cytokinesis. We show that one such factor is the kinesin KIF4A, which is present along the chromosome axes throughout mitosis and the central spindle in anaphase. These two pools of KIF4A depend on condensin I and PRC1, respectively. Previous work has shown KIF4A is activated by Aurora B at the anaphase central spindle. However, whether or not chromosome-associated KIF4A bound to condensin I is regulated by Aurora kinases remain unclear. To determine the roles of the two different pools of KIF4A, we generated specific point mutants that are unable to interact with either condensin I or PRC1, or are deficient for Aurora kinase regulation. By analysing these mutants, we show that Aurora kinases phosphorylate the condensin I dependent pool of KIF4A and thus actively promote chromosome congression from the spindle poles to the metaphase plate.

## Introduction

In prometaphase and metaphase, the two Aurora kinases A and B regulate chromosome congression by monitoring the process of chromosome bi-orientation (Lampson and Cheeseman, 2011). Tension is generated across the kinetochores of bi-oriented chromosomes enabling these to be discriminated from other attachment geometries by an error correction pathway. This process is primarily regulated by centromeric Aurora B which is physically close enough to the outer kinetochore to exert its effects only in the absence of tension, and thus provides a molecular read-out of bi-orientation and tension (Lampson and Cheeseman, 2011; Wang et al., 2011). Phosphorylation of microtubule-binding proteins of the outer kinetochore by Aurora kinases results in weakened interactions with incoming microtubules and thus facilitates the resolution of incorrect attachments resulting in error correction (DeLuca et al., 2011; Welburn et al., 2010). Uncongressed chromosomes close to the poles of the mitotic spindle come in to proximity of Aurora A. Aurora A, like Aurora B, can phosphorylate sites in the outer kinetochore implicated in microtubule binding, and thus promote release of chromosomes from the mitotic spindle (DeLuca et al., 2018; Ye et al., 2015).

Aurora kinases also promote chromosome bi-orientation and spindle bipolar assembly by inhibiting the activity of kinesin motor proteins. MCAK and KIF18B, which regulating the stability of microtubules attached to kinetochores are inhibited by Aurora A and B dependent phosphorylation (Andrews et al., 2004; Ems-McClung et al., 2013; McHugh et al., 2019; Tanenbaum et al., 2011; Zhang et al., 2008). Aurora A also phosphorylates and inhibits the centromere-associated kinesin CENP-E involved in efficient chromosome congression (Kapoor et al., 2006; Kim et al., 2010). A phosphorylation site mutant form of CENP-E cannot be inhibited by Aurora A and thus traps non-bioriented chromosomes at the spindle poles (Kapoor et al., 2006; Kim et al., 2010). Aurora A kinase is therefore thought to promote chromosome congression and error correction through inhibitory mechanisms that prevent trapping of chromosomes at the poles of the mitotic spindle.

This simple model poses a problem, since non-equatorial chromosomes will be released from bound microtubules by either the action of Aurora A or B, and thus fail to undergo movement towards the cell equator. How is congression of these non-equatorial chromosomes actively promoted? One possibility is that Aurora kinases, in addition to inhibitory actions, also facilitate chromosome congression by modulating the strength of the polar ejection force generated by chromokinesins.

Chromokinesins are a small subgroup of the kinesin superfamily which have binding sites for both microtubules and DNA (Almeida and Maiato, 2018; Mazumdar and Misteli, 2005). The ability to bind both to chromosomes and microtubules is thought to enable chromokinesins to generate the polar ejection force. This aids chromosome congression to the metaphase plate by pushing the chromosome and chromosome arms away from the poles and towards the cell equator (Rieder and Salmon, 1994). Chromokinesins KID and KIF4A of the kinesin-10 and kinesin-4 families, respectively, are the best studied. Initially, KID was thought to be the major kinesin creating the polar ejection force in human cells (Brouhard and Hunt, 2005; Levesque and Compton, 2001). However, subsequent studies showed that effective chromosome congression in cells requires cooperation of KID with KIF4A (Stumpff et al., 2012; Wandke et al., 2012).

KIF4A is particularly interesting in this context because it is known to be regulated by Aurora B (Bastos et al., 2014; Nunes Bastos et al., 2013). Early in mitosis, KIF4A plays an important role in both chromosome alignment and chromosome condensation (Mazumdar et al., 2004; Samejima et al., 2012; Stumpff et al., 2012; Takahashi et al., 2016; Wandke et al., 2012). In anaphase, KIF4A regulates microtubule dynamics and central spindle length (Kurasawa et al., 2004; Zhu and Jiang, 2005). These different functions are reflected in the different localizations of KIF4A. Throughout mitosis KIF4A localises to chromatin, while in anaphase a pool of KIF4A re-locates to the spindle midzone (Mazumdar et al., 2004). The temporally and spatially distinct functions of KIF4A are mediated by specific interaction partners at these different sites: the structural maintenance of chromosomes (SMC) protein complex condensin I on chromatin (Takahashi et al., 2016), and the microtubule binding and bundling protein regulator of cytokinesis 1 (PRC1) at the central spindle, respectively (Gruneberg et al., 2006; Kurasawa et al., 2004; Zhu and Jiang, 2005).

Condensin complexes localise to the axes of chromosomes in mitosis, and are essential for chromatin compaction and chromosome organisation (Hirano, 2012). Mammalian cells have two distinct condensin complexes, condensin I and II, which share the SMC subunits SMC2 and SMC4 but differ in the composition of their non-SMC subunits, CAP-D, G and H (Ono et al., 2003). KIF4A localises to chromosome arms and interacts specifically with the condensin I complex (Takahashi et al., 2016). In the absence of KIF4A, chromosomes become over condensed and shortened along their axes (Samejima et al., 2012; Takahashi et al., 2016). In anaphase, a second pool of KIF4A interacts with PRC1, a highly conserved, homo-dimeric microtubule binding protein required for central spindle formation (Mollinari et al., 2002; Mollinari et al., 2005). PRC1 and KIF4A are mutually dependent on each other for localisation to the anaphase spindle midzone where they together determine central spindle length (Bieling et al., 2010; Subramanian et al., 2010). This is achieved by molecular tagging of the ends of microtubules in the central spindle zone by KIF4A and PRC1 in a microtubule-length dependent manner (Subramanian et al., 2013). Phosphorylation of KIF4A at T799 close to the motor domain by the mitotic kinase Aurora B stimulates the motor activity of KIF4A, and is necessary for the anaphase function of KIF4A (Nunes Bastos et al., 2013).

While it is clear that condensin I and PRC1 contribute to the differential localisation of KIF4A to distinct cellular structures, the specific signal in KIF4A that mediates the interaction with PRC1 has not been identified. Furthermore, it is unclear whether activation of the KIF4A motor activity by Aurora kinases is relevant for the function of KIF4A and condensin I in chromosome condensation and congression. The interaction of KIF4A with condensin I is not directly regulated by Aurora B (Poonperm et al., 2017), eliminating this simple possibility. To understand the function of KIF4A at chromatin, we therefore identified and expanded the motifs in KIF4A required for interaction with condensin I and PRC1, respectively. This enabled us to analyse of the different aspects of KIF4A functionality in isolation for the first time, and test the idea that Aurora kinases contribute to chromosome congression by stimulating the motor activity of the chromosome bound pool of the chromokinesin KIF4A.

## Results

### Isolation of KIF4A mutants disrupted for either condensin I or PRC1 binding

In anaphase, KIF4A is present midway between the segregating chromosomes in a region defined by the anaphase spindle midzone protein PRC1 (Figure 1A). In both metaphase and anaphase, a discrete population of KIF4A concentrates on the axes of chromosomes similar to the condensin subunit SMC2 (Figure 1B). To understand the specific functions of KIF4A at the chromosome axis and central spindle, we searched for specific point mutants in KIF4A that would specifically disrupt the interactions with either condensin I or PRC1. Alignment of the C-terminal tail of metazoan KIF4A reveals the presence of a series of conserved motifs (Figure 2A). The conserved C-terminal region of KIF4A containing the cysteine rich domain (CRD) has previously been implicated in chromatin binding (Wu and Chen, 2008). In addition, the IXE motif SPIEEE has been implicated in binding to the phosphatase PP2A-B56 (Hertz et al., 2016). Between these two regions, are three absolutely conserved sequences. Two diphenylalanine clusters, FF1154 and FF1220, flank a positively charged lysine/arginine basic patch. The second of these motifs, FF1220 is needed for condensin I binding (Takahashi et al., 2016). Because diphenylalanine associated with a charge cluster has been implicated in other protein-protein interactions (Kaiser et al., 2005; Loewen et al., 2003), we focussed on this region in our search for the PRC1-binding motif.

**Figure 1.**
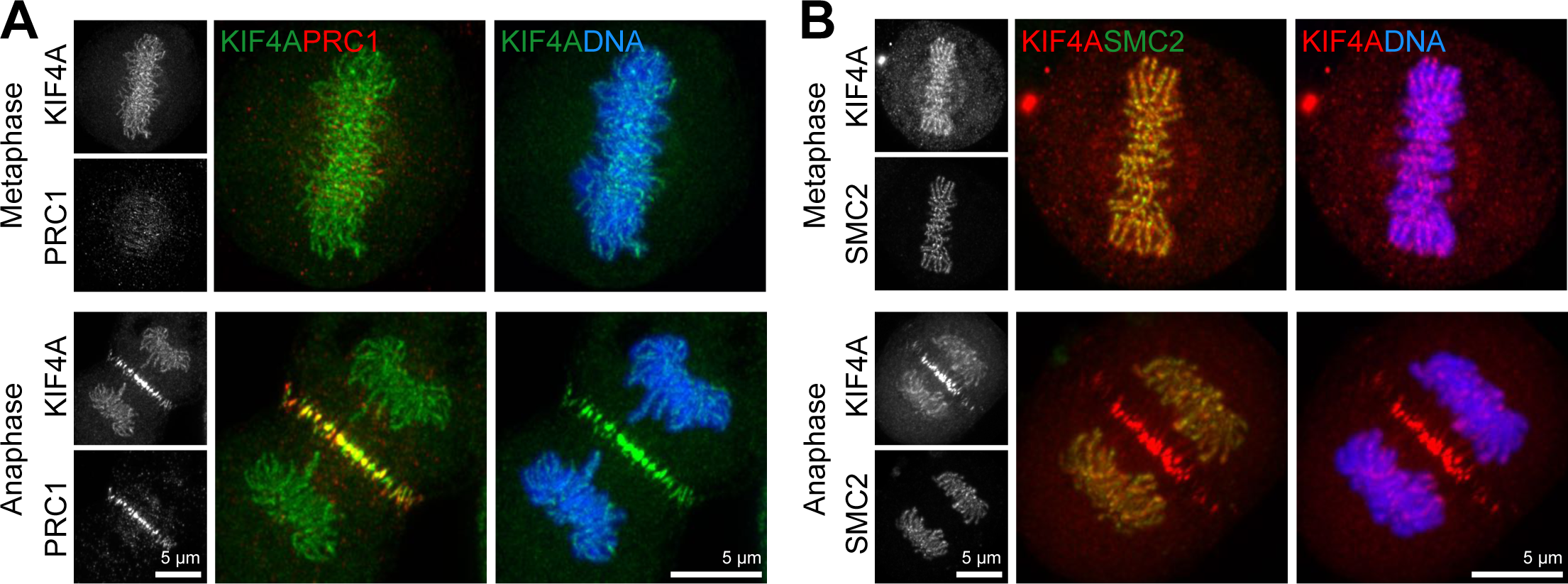
KIF4A localises to chromosomes axes and the anaphase central spindle. (A) Super-resolution images of eGFP-KIF4A and PRC1 or (B) SMC2-GFP and mScarlet-KIF4A were acquired using a Zeiss LSM880 microscope with Airyscan. DNA was detected using Hoechst 33258. Metaphase and anaphase localisations are shown. Scale bars mark 5µm.

**Figure 2.**
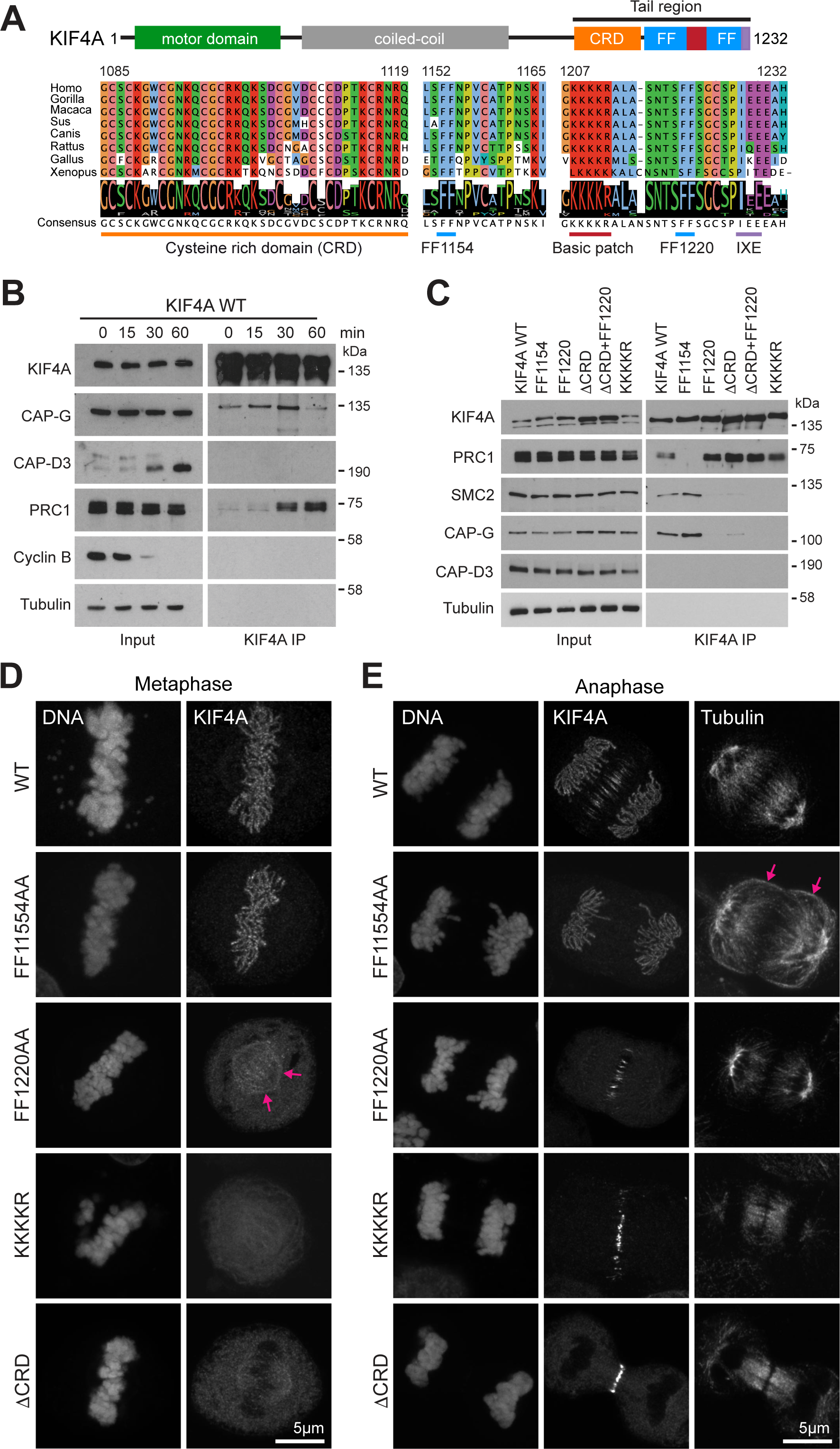
Two conserved FF-motifs in the C-terminal tail of KIF4A support interaction with either condensin I or PRC1. (A) Sequence alignment of KIF4A C-terminal tail in a group of metazoans showing a high degree of conservation. (B) Endogenous KIF4A was immunoprecipitated from HeLa cells exiting mitosis. A 0min time point was taken 25min after nocodazole wash-out. Synchronous progression of mitotic cells into anaphase was triggered with 200µM MPS1 inhibitor (AZ3146) to override the spindle checkpoint. The temporal association of KIF4A to Condensin I and PRC1 was followed over time by western blot. (C) Immunoprecipitation of wild type Flag-KIF4A (WT) and alanine point mutants identified through the sequence alignment from anaphase HeLa cells. (D) Super resolution images of FlpIn/TREx eGFP-KIF4A Hela induced with doxycycline to express KIF4A WT and point mutants in metaphase and (E) anaphase. Arrows mark the weak metaphase spindle staining for KIF4A FF1220AA, and the cortical spread of microtubules in the FF1154AA PRC1-binding mutant in anaphase.

To test the role of these conserved features they were either deleted or mutated to alanine and the corresponding KIF4A complexes isolated by immunoprecipitation from Hela cells in anaphase, when both the PRC1 and condensin I interactions can be detected (Figure 2B). This revealed that the KIF4A FF1154AA mutant fails to interact with PRC1 but retains its ability to bind condensin I detected using CAP-G and SMC2 (Figure 2C). Conversely, the FF1220AA and basic patch mutants failed to bind condensin I but still interacted with PRC1 (Figure 2C). Deletion of the CRD reduced, but did not abolish, binding to condensin I and had no effect on PRC1 interaction.

### Condensin I and PRC1-binding mutants of KIF4A show specific localisation defects

We next explored the roles of the CRD, basic patch, FF1154 and FF1220 motifs on KIF4A localisation in metaphase and anaphase. For this purpose, Hela Flp-In T-Rex cell lines expressing inducible copies of wild type or mutant GFP-KIF4A were generated. In comparison to wild type GFP-KIF4A, the FF1154AA PRC1-binding defective mutant failed to localise to the anaphase spindle, but showed otherwise normal targeting to chromosome axes in both metaphase and anaphase (Figure 2D and 2E). The ΔCRD, basic patch and FF1220 mutants all failed to target to the chromosome axes in either metaphase or anaphase, but were present on the central spindle in anaphase (Figure 2D and 2E). These data support the view that targeting to the chromosome axis and central spindle are independent events mediated through different pathways (Takahashi et al., 2016). This conclusion is also consistent with the analysis of KIF4A complexes in cells exiting mitosis (Figure 2B and 2C), where the FF1154AA and FF1220AA, ΔCRD or basic patch mutants were defective for either PRC1 or condensin I binding, respectively.

Because temporal control of KIF4A is crucial for its function, the effects of the FF1154AA and FF1220AA mutations on KIF4A complexes during mitosis and mitotic exit were followed (Movies 1-4). This confirmed that the FF1154AA mutation is defective for PRC1 interaction but shows condensin I binding in metaphase and anaphase (Figure 3A–3C). In contrast, the FF1220AA mutant was defective for condensin I binding at all time points but still displayed anaphase binding to PRC1 (Figure 3A–3C). The diphenylalanine clusters at FF1154 and FF1220 thus define motifs required for interaction of KIF4A with PRC1 and condensin I, respectively. In addition to FF1220, the upstream CRD and basic patch regions are required for efficient condensin I binding, but appear to play no role in PRC1 binding (Figure 2C). Time lapse imaging showed that these mutants showed the localisation defects matching these binding defects. KIF4A FF1154 which cannot bind PRC1 was present only on chromatin, whereas the ΔCRD and FF1220 mutants localise only to the anaphase spindle (Figure 3D). These mutants therefore enable us to investigate the specific functions of KIF4A-condensin I and KIF4A-PRC1 complexes.

**Figure 3.**
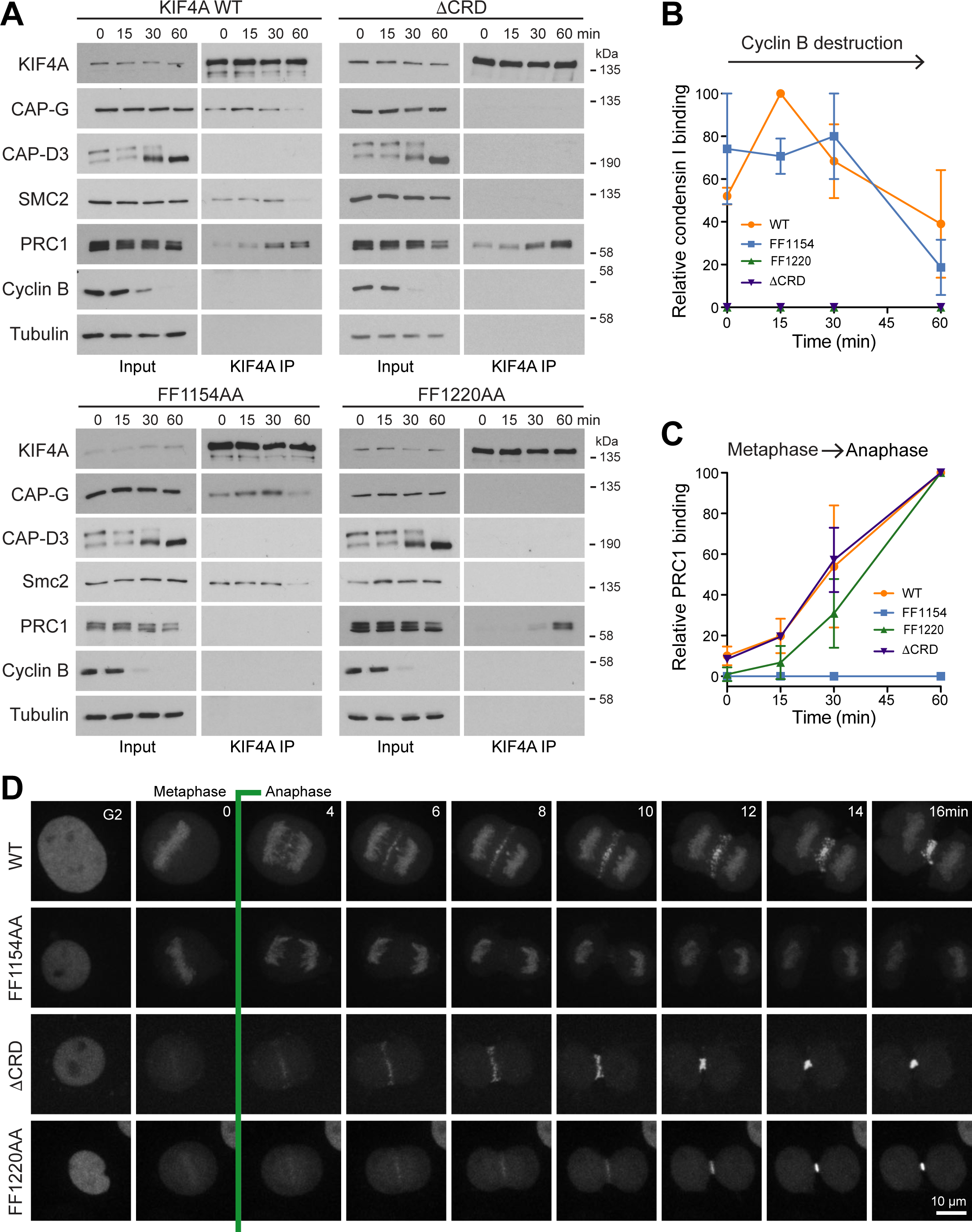
Temporal regulation of FF1220 condensin I and FF1154 PRC1-binding mutants of KIF4A. (A) KIF4A WT and mutant complexes were isolated from cells progressing from metaphase into anaphase. The temporal association of KIF4A to Condensin I and II and PRC1 was followed over time using western blot. (B) Relative KIF4A association to condensin I and (C) PRC1 was measured as a function of time and the mean plotted in the line graphs. Errors bars indicate the SEM for n=2. (D) HeLa cells depleted of endogenous KIF4A and expressing fluorescent protein tagged KIF4A WT, FF1154AA, ΔCRD or FF1220AA were imaged undergoing mitosis. Representative maximum projections are shown.

### PRC1-binding mutants of KIF4A are defective for central spindle length control

KIF4A is crucial for anaphase spindle length control and the formation of organised bundles of anaphase spindle microtubules (Figure S1A). When it is depleted, PRC1, components of the MKLP1-centralspindlin complex and cytokinesis factors such as ECT2 spread out along microtubules, rather than exhibiting a focussed localisation to anti-parallel microtubule overlap regions at the cell equator (Figure S1B). To test the specific role of the KIF4A-PRC1 interaction at the anaphase spindle, wild type and mutant GFP-KIF4A were expressed in HeLa cells depleted of endogenous KIF4A with siRNA directed to the 3’-UTR of the KIF4A mRNA. First, the distribution of PRC1 was examined to test if the loss of binding to KIF4A had an effect on its localisation in anaphase. This revealed that the PRC1 signal is spread over a broader area of the anaphase spindle in cells expressing the FF1154AA mutant when compared to wild type KIF4A or the condensin I binding defective FF1220AA and ΔCRD KIF4A mutants (Figure 4A and 4B). Anaphase spindle length was also increased in the PRC1 binding defective FF1154AA mutant compared to wild type KIF4A or the condensin I binding defective mutants (Figure 4C and 3D). A more detailed analysis of the 3-dimensional distribution of PRC1 and the MKLP1-centralspindlin complex showed that the defined clustered microtubule bundles seen with wild type KIF4A are disorganised in the FF1154AA mutant (Figure 4D and S1C). This becomes most obvious when viewed along the long axis of the spindle using a 90° rotation of the image data. The diameter of the central spindle increases and the density of PRC1 and MKLP1 is reduced in the central region compared to the cell cortex (Figure 4D and S1C). As a consequence, cytokinesis initiates earlier in anaphase and cleavage furrows are broader and less symmetric in KIF4A FF1154AA compared to the control cells (Figure S1D). These data support the view that the KIF4A is required for regulation of anaphase spindle elongation through its interaction with PRC1, rather than acting as a global regulator of microtubule dynamics.

**Figure 4.**
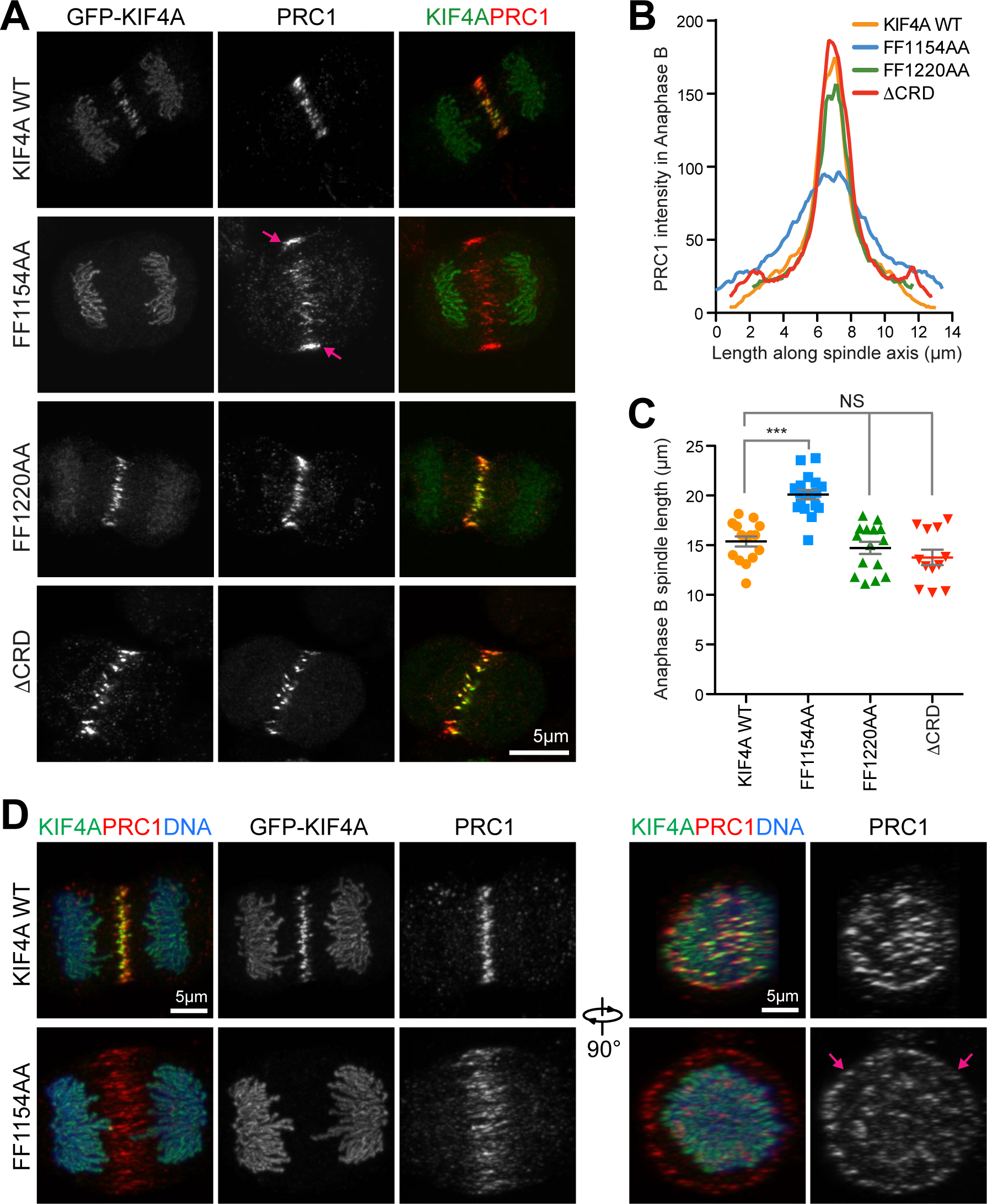
Defective anaphase spindle structure in the FF1154 PRC1-binding mutant of KIF4A. (A) Super resolution images of wild type KIF4A and mutants, showing anaphase localisation of KIF4A and PRC1. Arrows mark the spread of PRC1 signal at the central spindle. (B) The distribution of detected endogenous PRC1 staining throughout the length of the spindle in coverslips treated with siKIF4A and rescued with KIF4A WT or specific point mutants. Curves were centred to match the maximum intensity point (WT n=12, FF1154AA n=16, FF1220AA n=11, ΔCRD n=7). (C) The scatter plot represents the maximum length reached by the spindle in anaphase B and was obtained measuring data acquired by live cell imaging of HeLa cells treated with KIF4A siRNA and rescue with KIF4A WT and mutant constructs. The graph shows the mean with SEM (WT n=15, FF1154AA n=18, FF1220AA n=15, ΔCRD n=12). The nonparametric Kruskal-Wallis test was performed (P<0.0001) and conditions were compared post analysis by Dunns test (***, P<0.001). See also Figure S1 and Supplementary movies. (D) Super resolution images of KIF4A WT and the FF1154AA mutant, showing anaphase localisation of KIF4A and PRC1. Stacks were acquired to capture the total volume on the cell and the radial distribution of PRC1 at the cell equator is showed in the 90° rotation.

### Condensin I binding mutants of KIF4A are defective for chromosome morphology

The role of KIF4A in chromosome morphology was then investigated. As previously reported (Samejima et al., 2012), depletion of KIF4A results in significantly increased axial compaction of chromosomes in chromosome spread preparations (Figure 5A and 5B). This effect was seen in both HeLa and hTERT-RPE1 cells (S2A and S2B), and thus appears to reflect a general function for KIF4A in chromosome structure. The dependency of this function on the interaction with condensin I, PRC1 and requirement for Aurora-dependent phosphorylation was then examined. For this purpose, wild type and mutant GFP-KIF4A were expressed in Hela cells depleted of endogenous KIF4A with siRNA directed to the 3’-UTR of the KIF4A mRNA (Figure S2C). In this assay, wild type KIF4A and the PRC1 binding defective FF1154AA mutant rescued the axial compaction of chromosomes (Figure 5C and 5D). However, chromosomes in the condensin I binding defective FF1220AA mutant showed significantly increased axial compaction (Figure 5C and 5D), in agreement with previous work (Takahashi et al., 2016). The KIF4A-condensin I interaction is therefore important for normal chromosome structure. Importantly, the T799A/S801A phosphorylation mutant deficient for Aurora-regulated kinesin motor activity rescued axial compaction in this assay (Figure 5C and 5D). This agrees with previous work showing that Aurora kinases do not regulate the KIF4A-condensin I interaction (Poonperm et al., 2017), and also suggests that the function of KIF4A in chromosomes structure does not require Aurora-regulated kinesin motor activity. KIF4A therefore has discrete roles at chromosomes and the anaphase central spindle dependent on its specific binding partners at these sites.

**Figure 5.**
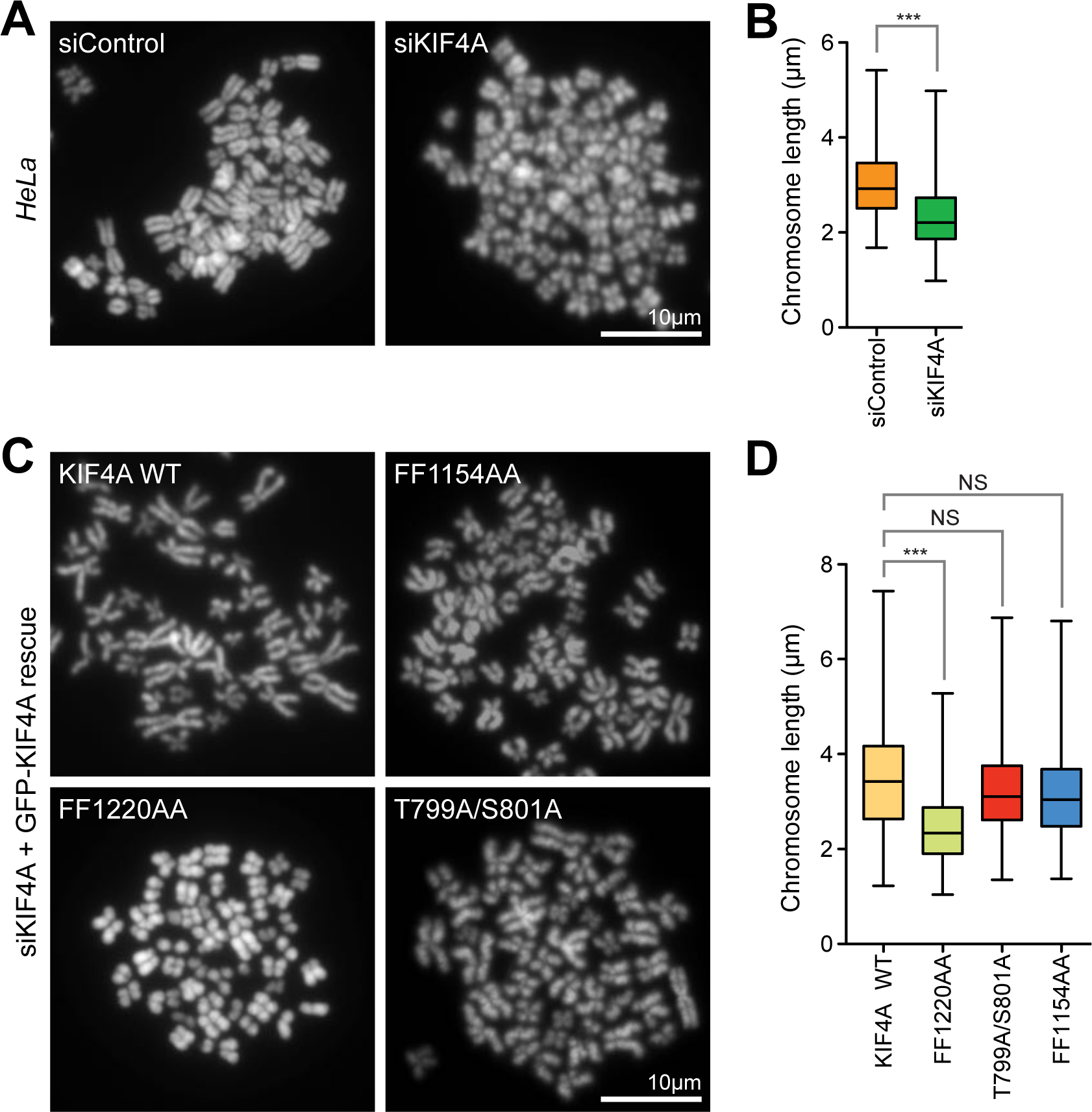
Chromosome condensation requires the KIF4A-condensin I interaction but not Aurora-dependent phosphorylation or PRC1 binding. (A) Chromosome spreads of HeLa cells transfected with control or siKIF4A. (B) Box and whiskers graph analyses chromosome length measured in FIJI. The analysis was done using an unpaired t test with Welch’s correction and 99% confidence intervals. P<0.001 ***. Whiskers show min and max. (siControl n=368, siKIF4A n=425). (C) Chromosome spreads of HeLa cells treated with siKIF4A and transfected with KIF4A WT and the indicted mutants. (D) Box and whiskers graph analyses chromosome length measured in FIJI. The analysis was performed with a nonparametric Kruskal-Wallis test (P<0.0001) and conditions were compared post analysis by Dunns test (***, P<0.001) (wild type KIF4A n=515, FF1220AA n=492, T799A/801A=653, FF1154AA n=660).

### Aurora regulation of the condensin-dependent pool of KIF4A promotes chromosome alignment

KIF4A has also been implicated in chromosome movement and the generation of the polar ejection force promoting congression of non-equatorial chromosomes to the metaphase plate (Stumpff et al., 2012; Wandke et al., 2012). This role requires the second chromokinesin KID, albeit the precise role of each kinesin remains unclear. Differences in their functions are expected since KIF4A and KID localise to different regions of the chromosome. KIF4A is present along the chromosome axis and KID shows a diffuse distribution across the chromosome (Figure 6A). To follow the role of KIF4A in chromosome congression, cells expressing a GFP-tagged version of the condensin subunit SMC2-GFP were used, since this enables individual chromosome arms to be clearly visualised. Depletion of both KIF4A and KID gave rise to chromosome alignment defects and delayed progression through mitosis (Figure 6B, arrows; S3A and S3B; Movie 5). Time lapse imaging show that most chromosomes were aligned at the metaphase plate with normal kinetics after 30min, but a small population remained at the spindle poles and showed delayed congression into the metaphase plate (Figure S3A, arrows). Furthermore, the movement of the chromosomes arms into the plate was delayed and only completed after ∼50-60min of mitosis. The ability of wild type or mutant KIF4A forms to rescue these defects was then tested (Movies 6-8). As expected, the PRC1 binding defective FF1154AA mutant supported chromosome congression and was indistinguishable from wild type KIF4A (Figure S3C), and was therefore not examined further. By contrast, the condensin I binding defective FF1220AA mutant failed to rescue chromosome congression (Figure 6C–6D and S3C) and showed significantly delayed progression into anaphase (Figure 6E). The T799A/S801A phosphorylation mutant also failed to rescue chromosome congression (Figure 6C–6D & S3C) and showed delayed progression into anaphase (Figure 6E). For both the FF1220AA and T799A/S801A mutants, compaction of chromosome arms into the metaphase plates was delayed until 50-60min of mitosis (Figure 6C and S3C). Chromosome congression therefore requires both recruitment of KIF4A to the axes of chromosomes by condensin I and Aurora-regulation of kinesin motor activity. Notably, this is different to the role in chromosome condensation which is supported by the T799A/S801A phosphorylation mutant and thus does not require Aurora-regulated kinesin motor activity.

**Figure 6.**
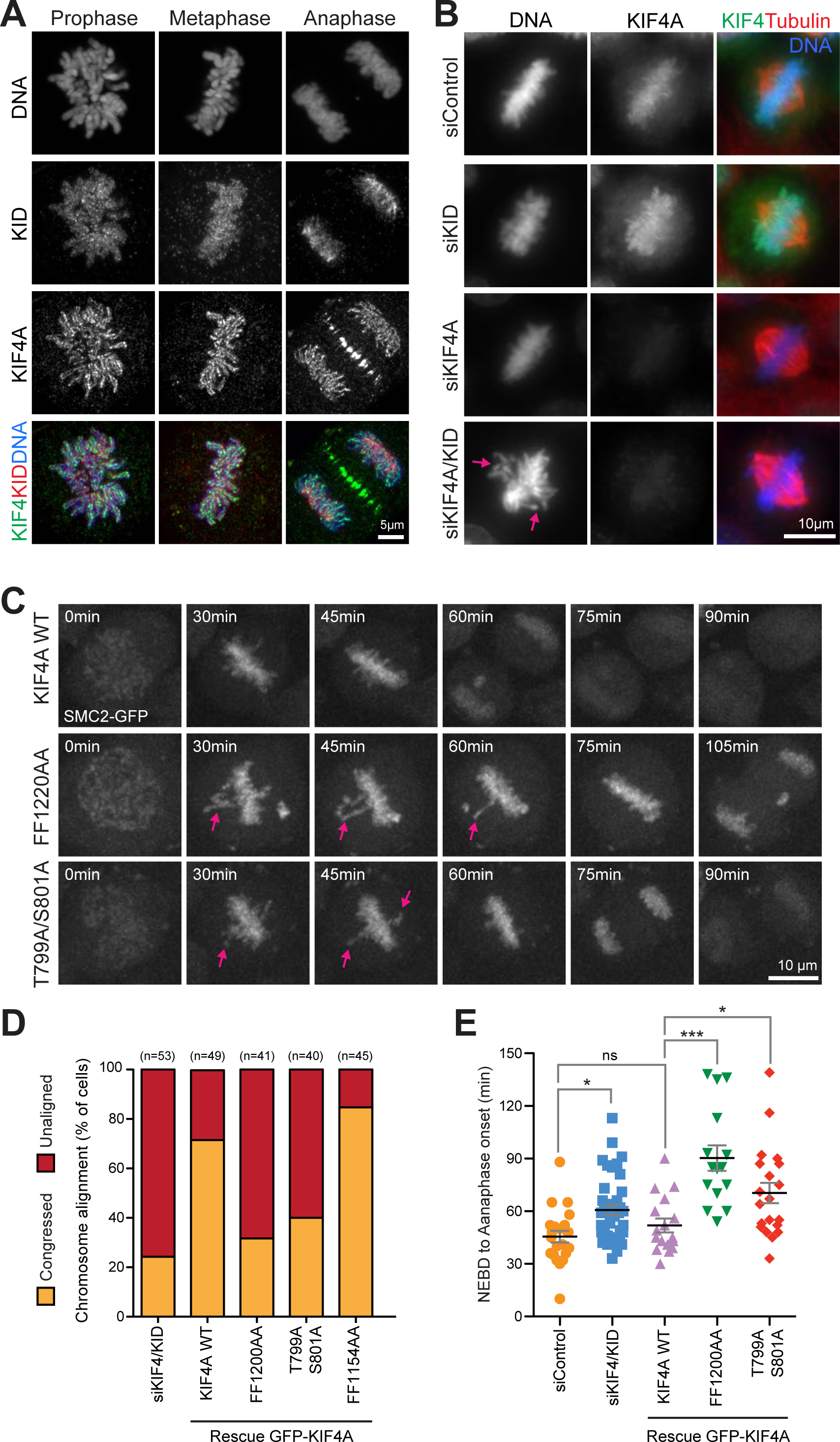
KIF4A but not the T799/S801 phosphorylation site mutant promotes prometaphase chromosome congression. (A) Localisation of KIF4A and KID in mitosis. CRISPR tagged eGFP-KIF4A cells were fixed and stained for endogenous KID. Images were acquired with an Airyscan system. (B) HeLa cells were treated for 48h before thymidine synchronisation with control, siKIF4A, siKID and siKIF4A+KID. After thymidine release and fixation, cells were stained for endogenous KIF4A. (C) mScarlet-KIF4A WT, FF1220AA or T799A/S801A were expressed in a CRISPR edited SMC2-eGFP cell line co-depleted for KIF4A and KID. Cells were imaged undergoing mitosis and representative maximum projections of the SMC2-eGFP signal are shown. Arrows mark chromosomes and chromosome arms showing delayed congression to the metaphase plate. (D) A column graph displays the extent of chromosome alignment for control and KIF4A and KID depleted cells and for KIF4A WT and mutants. (E) The scatter plot measures the mean time from NEBD to anaphase onset in SMC2-eGFP cells transfected with either siControl or siKIF4A+KID and rescued with KIF4A WT and mutants (siControl n=22, siKIF4A+KID n=59, rescue wild type KIF4A n=16, FF1220AA n=14, T799A/S801A n=20). The analysis was performed with a nonparametric Kruskal-Wallis test (P<0.0001) with a 95% confidence intervals and conditions were compared post analysis by Dunns test (***, P<0.001; *, P<0.1).

Aurora A and B create defined non-overlapping phosphorylation gradients within the spindle, at the spindle poles and centromeres, respectively (Figure 7A). To assess the contributions Aurora A and B make to KIF4A regulation in chromosome alignment, their role in generating the T799/S801 phosphorylation in prometaphase was examined using specific antibodies. Consistent with a role for Aurora-dependent phosphorylation of KIF4A both in early and late mitosis, T799 phospho-antibodies (pT799) show that KIF4A phosphorylation rises as cell enter mitosis with a similar profile to the T288 activating phosphorylation on Aurora A (pT288) and the S10 phosphorylation of Histone H3 (pS10) created by Aurora B (Figure S4A). This is rise in pT799 is slightly delayed when compared to CDK-dependent phosphorylation detected using PRC1 pT481, and persists longer into anaphase (Figure S4B, 12-13h time points). In cells with unaligned chromosomes, KIF4A pT799 is detected at regions overlapping with both Aurora A and B at the spindle poles and centromeres, respectively (Figure 7B). These signals are lost when KIF4A is depleted, leaving only a non-specific centrosome signal at each spindle pole (Figure 7B). KIF4A has been previously shown to be a target of Aurora B (Nunes Bastos et al., 2013), and we therefore tested if it can also be phosphorylated by Aurora A. In vitro kinase assays showed that recombinant KIF4A can be phosphorylated by Aurora A and that this activity is inhibited by the specific Aurora A inhibitor MLN8537 (Figure 7C). We then investigated the contribution of Aurora A to KIF4A phosphorylation in cells.

**Figure 7.**
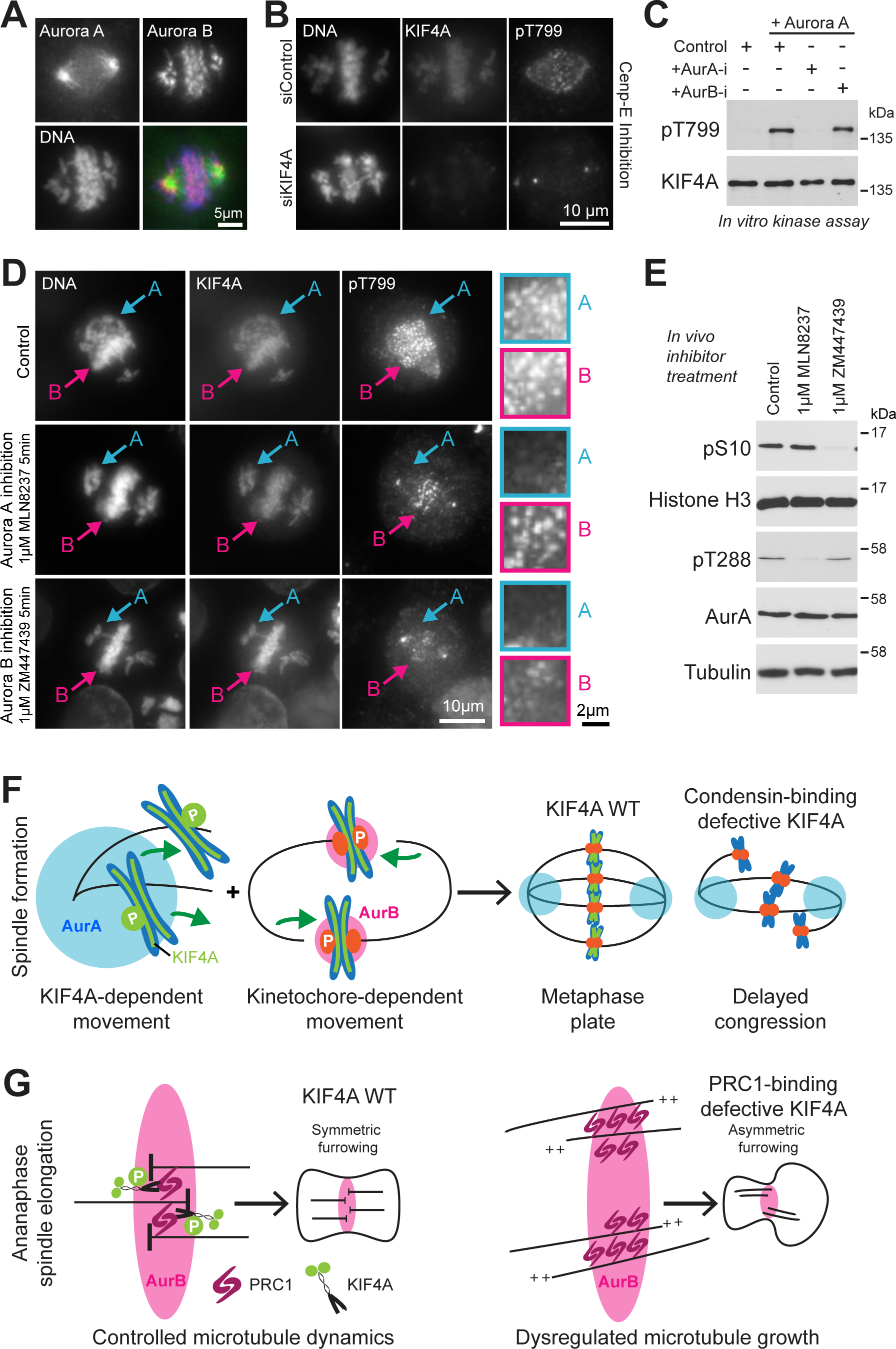
Aurora A phosphorylates KIF4A on off-axis chromosomes. (A) At 7h after thymidine release, HeLa cells were treated with 300nM of CENP-E inhibitor (GSK923295) for 3h to accumulate chromosomes at the spindle poles. The cells were then stained for DNA, Aurora A and Aurora B. (B) CRISPR edited eGFP-KIF4A cells treated either with siControl or siKIF4A and CENP-E inhibitor were stained for KIF4A pT799 and DNA. (D) KIF4A pT799 staining following 5min treatment with DMSO (Control), 1µM Aurora A inhibitor (AurA-I; MLN8237) or 1 µM Aurora B inhibitor (AurB-I; ZM447439). Arrows A and B mark the phosphorylation signals of KIF4A on chromosomes in the Aurora A and B regions of the spindle, respectively. The enlargements on the right show the pT799 signal in the same regions. (E) Western blots show the specificity of the Aurora A and B inhibitors under these conditions. (F) The model summarises Aurora-regulated kinetochore-dependent and independent pathways contributing to chromosome congression. At kinetochores, Aurora B destabilises kinetochore-microtubule attachments for non-bioriented chromosomes. Aurora A phosphorylates the condensin I bound pool of KIF4A at polar chromosomes, promoting KIF4A motor activity and chromosome congression. (G) In anaphase, PRC1 and KIF4A control microtubule growth and the organisation of the anaphase spindle, ensuring symmetric furrowing. The KIF4A mutant FF1154AA shows this interaction is essential for regulated microtubule growth and organised cytokinesis.

In control cells, unaligned chromosomes close to the Aurora A polar region or undergoing alignment at the equator show punctate pT799 signals (Figure 7D). When Aurora A was inhibited, the pT799 signal on the polar, non-equatorial chromosomes was lost within 5 minutes (Figure 7D, Aurora A inhibition – A) while the signal on equatorial chromosomes was retained (Figure 7D, Aurora A inhibition – B). Inhibition of Aurora B resulted in loss of the equatorial signal for pT799 (Figure 7D, Aurora B inhibition – B). Interestingly, Aurora B inhibition also reduced the signal for pT799 on chromosomes closer to the poles, suggesting there is interplay between Aurora A and B away from the metaphase plate. Western blot shows that in cells the two inhibitors display the expected specificity for Aurora A (pT288) and Aurora B (pS10), respectively (Figure 7E). In summary, we conclude that the pT799 signal reflects the overlap between axial distribution of KIF4A and the two kinases. Aurora A and B can therefore both phosphorylate chromosome-associated KIF4A, and we conclude stimulate its kinesin motor activity.

## Discussion

This work provides mechanistic insight by explaining how Aurora-kinase gradients within the spindle promote chromosome congression. We show that Aurora-dependent regulation of the pool of KIF4A recruited to the chromosome axes through interaction with condensin I is important for chromosome congression into a compact metaphase plate. This role is different to the function in chromosome structure, which also requires interaction with condensin I but does not require the phosphorylation-stimulated kinesin motor activity. Aurora kinases therefore play a crucial role in activating KIF4A function during chromosome movements and thus complement the kinetochore and CENP-E dependent capture and bi-orientation pathway (Figure 7F). Previous work has explained how Aurora A and B uncouple chromosomes from microtubules by phosphorylation of outer kinetochore proteins and the kinesin motor CENP-E (DeLuca et al., 2011; DeLuca et al., 2018; Kim et al., 2010; Welburn et al., 2010). We now propose that Aurora A phosphorylates and activates KIF4A, thus promoting the kinetochore-independent congression of chromosomes and chromosome arms from the cell poles into the metaphase plate (Figure 7F). Without this pathway, non-equatorial chromosomes would be released from bound microtubules by either the action of Aurora A or B, and thus fail to undergo movement towards the cell equator. This failure is seen in cells expressing the condensin I and Aurora-phosphorylation mutants of KIF4A. We therefore conclude that Aurora kinases facilitate chromosome congression by modulating the strength of the KIF4A-dependent component of the polar ejection force. Our data also demonstrate that the interaction of KIF4A is necessary for proper regulation of chromosome structure by condensin I and that this effect is independent of Aurora-regulated motor activity or the interaction with PRC1 required for microtubule organisation at the anaphase central spindle (Figure 7G). However, this leaves open the question of how KIF4A acts. One simple possibility is that the dimeric C-terminal tail of KIF4A which has two copies of FF1220 would act to link two condensin I complexes. Further work will be necessary to test this idea.

Here, we provide important new insights into the regulation of the polar ejection force by Aurora kinases. Nonetheless, a number of questions relating to this process remain to be addressed. KIF4A phosphorylation is rapidly lost within 5 minutes of Aurora A or B inhibition, and thus there must be a phosphatase counteracting these Aurora kinases. Previous reports show that PP2A-B56 interacts with KIF4A at the central spindle and dephosphorylates T799 in anaphase (Bastos et al., 2014; Hertz et al., 2016). However, KIF4A does not show co-localisation with PP2A-B56 along the chromosome axes early in mitosis. Additionally, the binding site mapping data, summarised in Figure 2A, indicate that the condensin I binding site is immediately adjacent to the PP2A-B56 binding motif and that binding of these two large multi-subunit complexes is thus likely to be mutually exclusive. This suggests that there is another phosphatase or pool of PP2A-B56 that acts on chromatin-bound KIF4A. One possibility is that this could be PP1, as in the case of CENP-E (Kim et al., 2010). Further questions relate to the regulation of KID and its precise mechanism of localisation to chromatin in prophase, prometaphase and anaphase. Answering these will be important to fully understand high fidelity capture and segregation of chromosomes by the mitotic spindle.

## Materials and Methods

### Reagents and antibodies

General laboratory chemicals and reagents were obtained from Sigma-Aldrich and Thermo-Fisher Scientific unless specifically indicated. Inhibitors for Aurora A (MLN8237, Cambridge Bioscience), Aurora B, (ZM447439, Tocris Bioscience), MPS1 (AZ3146, Tocris Bioscience), and CENP-E (GSK923295, Selleckchem) were dissolved in DMSO to make 10mM stocks. Thymidine (Sigma-Aldrich, 100mM stock) and doxycycline (Invivogen, 1mM stock) were dissolved in water. A 1mg/ml stock of the DNA dye Hoechst 33258 (Sigma) was dissolved in water.

Commercial antibodies were used to detect PRC1 pT481 (2189–1; Epitomics), cyclin B1 (05–373; EMD Millipore), tubulin (T6199; Sigma-Aldrich), FLAG epitope tag (F7425; Sigma-Aldrich). Smc2 (ab10399, Abcam), NCAP-G (A300-602A; Bethyl), NCAP-D3 (ab70349; Abcam), KID/KIF22 (ab222187; Abcam), Aurora A (4718S; Cell Signalling), Aurora B (mouse AIM1; Becton Dickinson), Aurora A pT288 (3079S; Cell Signalling), H3 (4499S; Cell Signalling), Histone H3 pS10 (6G3; Cell Signalling). Antibodies against GFP, mCherry, PRC1, ECT2, KIF23 (MKLP1), KIF4A and KIF4A pT799 have been described previously (Nunes Bastos et al., 2013). Secondary donkey antibodies against mouse, rabbit, or sheep and labelled with Alexa Fluor 488, Alexa Fluor 555, Alexa Fluor 647, Cy5, or HRP were purchased from Molecular Probes and Jackson ImmunoResearch Laboratories, Inc., respectively. Affinity purified primary and HRP-coupled secondary antibodies were used at 1µg/ml final concentration. For western blotting, proteins were separated by SDS-PAGE and transferred to nitrocellulose using a Trans-blot Turbo system (Bio-Rad). All western blots were revealed using ECL (GE Healthcare). Protein concentrations were measured by Bradford assay using Protein Assay Dye Reagent Concentrate (Bio-Rad).

### Molecular biology

All DNA primers were obtained from Invitrogen. Human KIF4A was amplified using Pfu hot start turbo polymerase (Agilent Technologies). Mammalian expression constructs were made in pcDNA5/FRT/TO (Invitrogen) modified to encode eGFP, mScarlet or FLAG tags. Mutagenesis was performed using the QuickChange method (Agilent Technologies). KIF4A siRNA 5’-CAGGTCCAGACTACTACTC-3’ against the 3’-UTR was obtained from QIAgen, and an optimised siRNA pool for KIF22 (KID) was obtained from Dharmacon (L-004962-00-0005). Control duplexes for siRNA have been described previously (Cundell et al., 2013).

### Cell culture procedures

HeLa cells and HEK293T were cultured in DMEM with 1% [vol/vol] GlutaMAX (Life Technologies) containing 10% [vol/vol] bovine calf serum at 37°C and 5% CO_2_. RPE1 cells were cultured in DME/F-12 (Sigma) supplemented with 1% [vol/vol] GlutaMAX (Invitrogen) and containing 10% [vol/vol] bovine calf serum at 37°C and 5% CO_2_. Mirus LT1 (Mirus Bio LLC) was used to transfect all cell lines. siRNA transfection was performed with Oligofectamine (Invitrogen) and TransIT-X2 (Mirus), for HeLa and RPE respectively. HeLa cell lines with single integrated copies of KIF4A wild type and the mutants described in the figures were created using the T-Rex doxycycline-inducible Flp-In system (Invitrogen). HeLa cell lines with GFP or mScarlet inserted in frame at the C-terminus of KIF4A and SMC2 in the endogenous loci were constructed using CRISPR/Cas9 (Alfonso-Perez et al., 2019).

### Isolation of KIF4A complexes and in vitro kinase assays

For each time course, four 15 cm dishes of HeLa cells, one dish per time point, were transfected with Flag-KIF4A constructs. After 24h, cells were arrested with 0.1µM nocodazole for 18h. Mitotic cells were collected by shake-off, pooling together dishes transfected with the same construct. Nocodazole was removed by washing three times with PBS, and then twice with growth medium pre-warmed to 37°C and equilibrated with CO_2_. Cells were then resuspended in 4.5ml growth medium and left in the incubator at 37°C for 25minutes to allow the reconstitution of the mitotic spindle. After harvesting a 0minute time point by washing 1ml of cells in ice cold PBS, 200µM AZ3146 (MPS1 inhibitor) was added to the remaining cell suspension and further samples taken at 15,30 and 60min. Cells were lysed on ice with 1ml of lysis buffer (20mM Tris-HCl pH7.4, 150mM NaCl, 1% (vol/vol) Igepal CA-630, 0.1% (wt/vol) sodium deoxycholate, 100nM okadaic acid, 40mM ß-glycerophosphate, 10mM NaF, 0.3mM Na_3_VO_4_, protease inhibitor cocktail (Sigma-Aldrich), phosphatase inhibitor cocktail (Sigma-Aldrich). supplemented with 40U of benzonase nuclease (E1014-Sigma) for 3mg of total lysate and 2mM of MgCl_2_. Flag-KIF4A complexes were isolated by 2h incubation at 4°C with 15µl anti-Flag M2-agarose beads (Sigma-Aldrich). Beads were washed twice with 1ml lysis buffer and twice with 1ml wash buffer (20mM Tris-HCl pH7.4, 150mM NaCl, 0.1% (vol/vol) Igepal CA-630, 40mM ß-glycerophosphate, 10mM NaF, 0.3mM Na_3_VO_4_) and resuspended in 50µl 3x Laemmli buffer for Western blotting. KIF4A in vitro kinase assays were performed using recombinant Aurora A (Aaronase) as described previously (Nunes Bastos et al., 2013).

### Fix cell imaging and super-resolution microscopy

For fixed cell imaging, cells were grown on No 1.5 glass coverslips (AGL46R16-15, Agar Scientific), washed once with PBS and then fixed in −20°C methanol for 5 minutes. Coverslips were washed 3 times with PBS and then stained. Primary and secondary antibody staining was performed in PBS for 60 minutes at room temperature. Coverslips were mounted in Moviol4-88 mounting medium (EMD Millipore). Images were acquired using a 60x NA1.35 oil immersion objective) on an upright microscope (BX61; Olympus) with filter sets for DAPI, GFP/Alexa Fluor 488, Alexa Fluor 555, Alexa Fluor 568, and Alexa Fluor 647 (Chroma Technology Corp.), an 2048×2048 pixel CMOS camera (PrimΣ; Photometrics), and MetaMorph 7.5 imaging software (Molecular Devices). Illumination was provided by an LED light source (pE300; CoolLED Illumination Systems). Image stacks of 10 to 17 planes with a spacing of 0.6µm through the cell volume were maximum intensity projected in MetaMorph.

For high-resolution imaging, doxycycline-inducible Flp-In/T-REx wild type or mutant eGFP-KIF4A HeLa cells were seeded on 22×22×0.17mm coverslips (AGL46S22-15) and transfected with siRNA against the 3’-UTR of endogenous KIF4A. After 48h, cells were synchronised by addition of thymidine to a final concentration of 2.5mM for 20h. Cells were then washed 3x in fresh growth medium, and expression of KIF4A then induced with 1µM doxycycline for 10h. Fixation was carried out by immersing the cells on coverslips in −20°C methanol for 5min, then washing 3x in 2ml room temperature PBS before staining with antibodies for tubulin and PRC1 as described in the figures. When the SMC2-GFP cell line was used, cells were transfected with mScarlet-KIF4A constructs and the signal was enhanced by staining with a sheep anti-mCherry primary antibody and Donkey anti-sheep Alexa568 secondary antibody. Tubulin and PRC1 were stained with antibodies. DNA was detected using DNA dye Hoechst 33258 diluted to 1µg/ml. In all cases, coverslips were mounted on slides using Vectashield (Fisher Scientific). Images were acquired using an inverted Zeiss LSM880 microscope fitted with an Airyscan detector and a Plan-Apochromat 63×/1.4-NA oil lens. The 488-nm Argon and 561-nm diode lasers in combination with a dual band emission filter (band pass 420-480nm and 495-550nm) were used either to excite GFP and mScarlet or to detect staining of tubulin and PRC1. DNA signal was detected using a solid-state laser. Sequential excitation of each wavelength was switched per Z-stack. Airyscan processing was performed using the ZEN system software. Z-stack maximum intensity projection was performed using FIJI (Schindelin et al., 2012) to produce TIFF images for the figures. Images were then cropped in Adobe Photoshop and placed into Illustrator to produce the figures. To visualize cells in 3D, an average of 100 stacks (0.14µm between Z-stacks) were acquired. The processed images were then opened in FIJI and a 3D projection with interpolation was performed. The 3D rendered images were saved both as “.avi” movie files and sequences of TIFF images for use in the figures.

### Live cell microscopy

For live-cell imaging, cells were plated in 35-mm dishes with a 14-mm No. 1.5 thickness coverglass window (MatTek Corporation) then treated as described in the figure legends. The dishes were placed in a 37°C and 5% CO_2_ environment chamber (Tokai Hit) on an inverted microscope (IX81; Olympus) with a 60X 1.42 NA oil-immersion objective coupled to an Ultraview Vox spinning disk confocal system running Volocity 6 (PerkinElmer). Images were captured with a EMCCD camera (C9100-13; Hamamatsu Photonics). Exposure time was 100 ms 4% 488-nm laser power for GFP and 6% 561-nm lasers for mScarlet. Typically, 15 planes, 0.6µm apart were imaged every minute. Spindle length measurements were performed on the original data with Volocity 6 software. Images for figures in TIFF format and movies were created in FIJI using maximum intensity projection of the fluorescent channels. Images were then placed into Adobe Illustrator CS5.

### Chromosome spreads

HeLa S3 and hTERT-RPE1 cells were seeded at low confluency (3 × 10^6^cells per 15 cm dish) then transfected with siControl or siKIF4A after 24h. After a further 24h cells were either left untransfected or transfected with eGFP-KIF4A constructs for 40h. At this stage, dishes were at 60-70% confluency and the cells were arrested with nocodazole 0.1 µm/ml for 6h. Mitotic cells were then collected by shake off and a small aliquot was saved to check depletion and overexpression levels. Cells were resuspended in 0.075 mM KCl pre-warmed at 37°C to allow cell swelling. After 20-30 minutes of incubation at 37°C, cells were pelleted at 1000 rpm for 5min and resuspended in the leftover volume of KCl. Fixative was prepared by mixing 3 volumes of methanol and one volume of acetic acid on ice. Fixation was performed adding gently 500µl of ice-cold fixative while vortexing at slow speed. Before performing the spread, super frost slides (AGL4341, Agar Scientific) were placed in cold water. Cells were resuspended in cold fixative and dropped onto a slide tilted at 45°C, washed gently with fixative solution and dried on a wet paper towel placed on top of a water bath set at 80°C. Coverslips were then mounted of the slides using Mowiol 4-88 containing 1µg/ml Hoechst 33258. Chromosomes were imaged using a 100x NA1.4 immersion oil objective taking a snapshot of a single plane. Images were then opened and analysed in FIJI and chromosomes length was measured manually.

## Supplemental Information

Figure S1 supports the analysis of the anaphase spindle and cytokinesis in Figure 4. Figure S2 shows chromosomes spreads for hTERT-RPE1 cells and extends Figure 5. Data on chromosome alignment is shown in Figure S3 to supplement Figure 6. Figure S4 shows analysis of KIF4A phosphorylation by Western blot to supplement Figure 7. Movies 1-4 show the localisation of wild type KIF4A or the FF1154AA, ΔCRD or FF1220AA mutants for Figure 3D. Movie 5 shows chromosome movement in control and siKIF4/KID cells and supports Figure S3A. Movies 6-8 show chromosome movement in KIF4A WT, FF1220AA or T799A/S801A expressing cells for Figure 6C.

## Acknowledgements

EP was supported by a Royal Society Newton Fellowship grant (NF161529) and FAB by a Cancer Research UK programme grant (C20079/A15940). RC was supported by a Medical Research Council Senior Non-Clinical Research fellowship awarded to UG (MR/K006703/1). We thank members of the Barr and Gruneberg labs for their advice during the work and comments on the manuscript.

The authors declare no competing financial interests.

## Author Contributions

Conceptualization: F.A. Barr. Investigation: E. Poser and R. Caous. Funding acquisition: F.A. Barr. Supervision: F.A. Barr and U. Gruneberg. Writing – original draft: F.A. Barr, U. Gruneberg and E. Poser. Writing – review and editing: E. Poser, R. Caous, U. Gruneberg and F.A. Barr.

