## Supplemental Figures for "Aurora A promotes chromosome congression by activating the condensin-dependent pool of KIF4A"

Please address correspondence to:

**Supplemental Figure Legends**

**Figure S1.** KIF4A/PRC1 defective binding mutant alters midzone formation and affect furrowing. A) Super resolution images of microtubule and spindle organisation in control and KIF4A depleted HeLa mCherry histone H2B/eGFP-tubulin cells . **(**B) Anaphase distribution of central spindle and cytokinesis regulators PRC1, MKLP1 and ECT2 in control and KIF4A depleted cells. (C) Super resolution images of wild type KIF4A and the FF1154AA mutant, showing the distribution of KIF4A and MKLP1 in anaphase. Stacks were acquired to capture the total volume on the cell and the radial distribution of MKLP1 at the cell equator is showed in the 90º rotation. (D) HeLa cells depleted of endogenous KIF4A and transient transfected with wild type eGFP-KIF4A WT or the FF1154AA mutant were imaged during mitosis. Representative maximum projections with a bright-field reference image are shown. Anaphase onset and furrowing are indicated. (E) Scatter plot measures the minutes from anaphase to onset of furrowing for WT n=19, FF1154AA n=22 and FF1220AA n=16. The nonparametric Kruskal-Wallis test was performed and conditions were compared post analysis by Dunns test with a 99% confidence intervals (***, P=0.001).

**Figure S2.** KIF4A regulates chromosome length in hTERT-RPE1 cells. A) Chromosome spreads of control and KIF4A depleted hTERT-RPE1 cells. (B) Box and whiskers graph analyses chromosome length measured in FIJI. The analysis was done using an unpaired t test with Welch's correction and 99% confidence intervals. P<0.0001. Whiskers show min and max. (siControl n=250, siKIF4A n=160). The blots show depletion of KIF4A when cells were treated with KIF4A siRNA. (C) Western blots showing depletion of endogenous KIF4A and expression of transfected eGFP-KIF4A constructs.

**Figure S3.** KIF4A and KID are required for chromosome congression. (A) Control, KIF4A, KID and KIF4A/KID depleted SMC2-eGFP cells were imaged during mitosis. Representative maximum projection of the GFP signal is shown. Arrows indicate non-congressed chromosomes and chromosome arms extending out of the metaphase plate. (B) Blots showing the depletion of either KIF4A, KID or both. (C) HeLa cells depleted of KIF4A and KID and transfected with eGFP-KIF4A constructs, were fixed and stained for DNA and tubulin.

**Figure S4.** KIF4A is phosphorylated in the early stages of mitosis and get dephosphorylated only in late anaphase. (A) HeLa cells were synchronised with a double thymidine block and followed for 15h after release. Cells were lysed directly in sample buffer every hour. Western blots were then performed for the proteins and phosphorylation sites indicated in the figure. (B) Relative intensity of KIF4A pT799, Histone H3 pS10, PRC1 pT481, Aurora A pT288 is plotted in the graph from 7 to 15h of the time course.

**Supplemental Movie Legends**

**Movie 1-4**

Related to Figure 3D. HeLa cells depleted of endogenous KIF4A were transfected with wild type KIF4A or the FF1154AA, ∆CRD or FF1220AA mutants. Cells were imaged 9 hours after thymidine release. One timepoint was captured every minute and the videos play at 7 fps. The eGFP and mScarlet signals are maximum intensity projections. A single plane reference was taken for the bright field.

**Movie 5**

Related to Figure S3A. Chromosome movement was imaged in SMC2-GFP cells treated with control siRNA or a pool of siKIF4A and siKID. Cells were imaged 9 hours after thymidine release. One timepoint was captured every minute and the videos play at 7 fps. The eGFP signal is a maximum intensity projection.

**Movies 6-8**

Related to Figure 6C. Chromosome movement was imaged in SMC2-GFP cells depleted of KIF4A and KID and then rescued by expression of wild type mScarlet-KIF4A, or the FF1220AA or T799A/S801A mutants. One timepoint was captured every minute and the videos play at 7 fps. The eGFP and mScarlet signals are maximum intensity projections.

**Figure S1.** Poser et al.

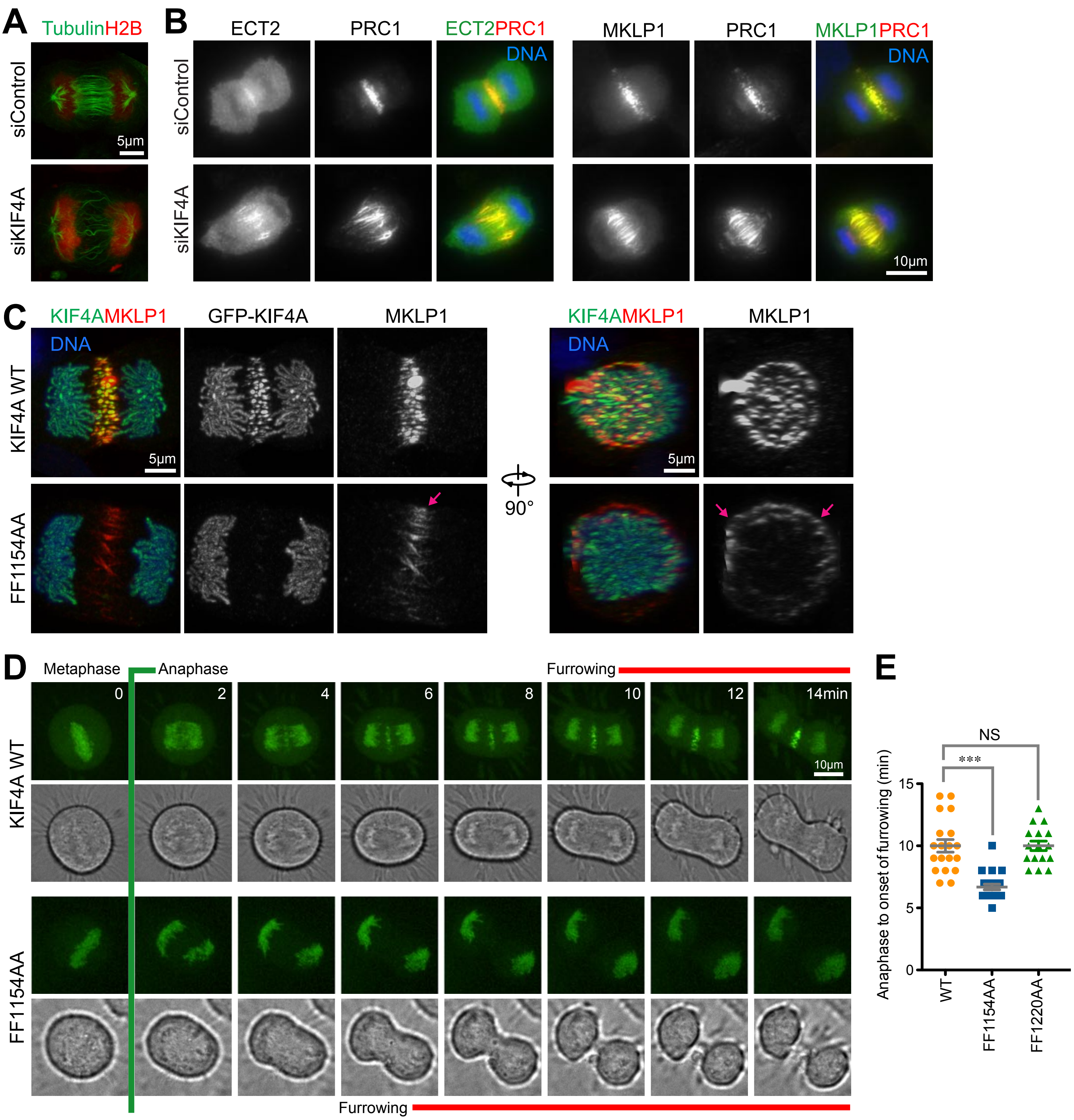

**Figure S2.** Poser et al.

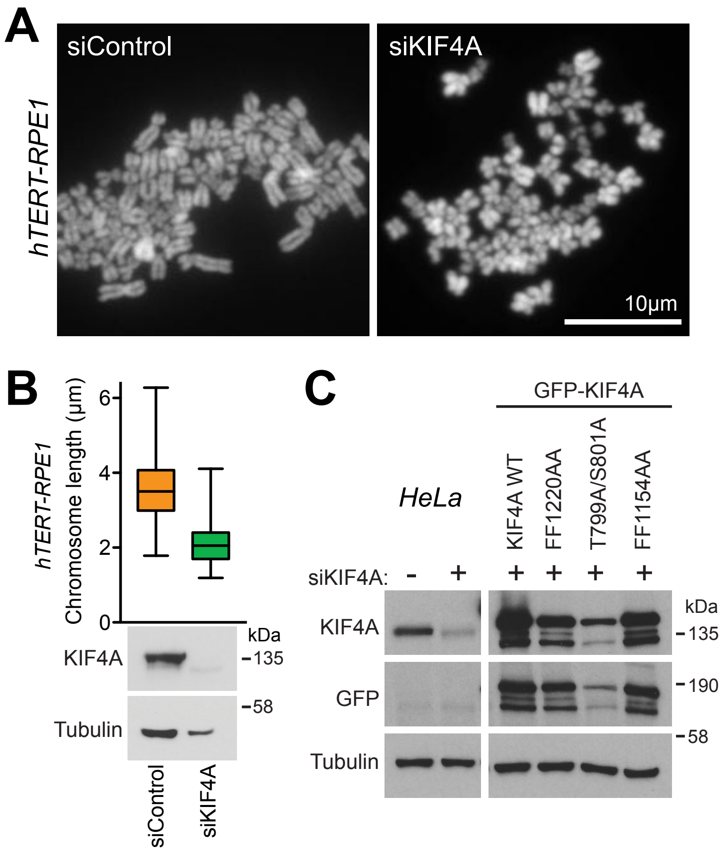

**Figure S3.** Poser et al.

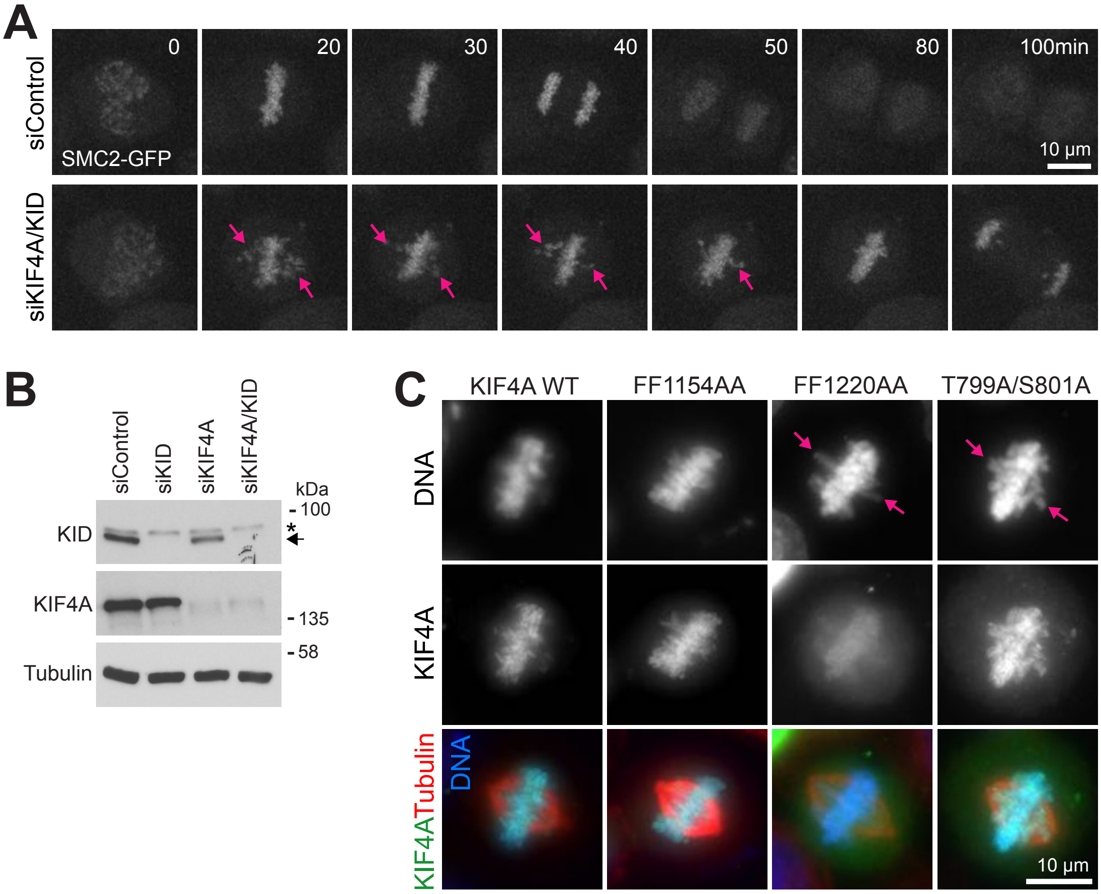

**Figure S4.** Poser et al.

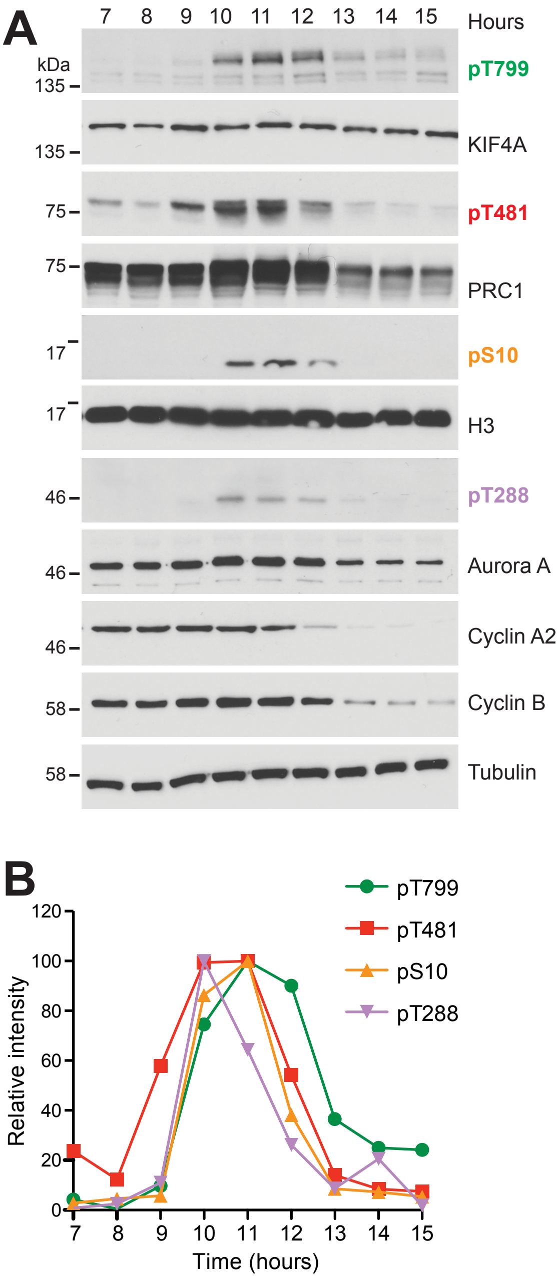
